# Lipid Tails Modulate Antimicrobial Peptide Membrane Incorporation and Activity

**DOI:** 10.1101/2021.08.12.456135

**Authors:** Lawrence R. Walker, Michael T. Marty

## Abstract

Antimicrobial peptides (AMPs) are cationic, amphipathic peptides that interact directly with lipid bilayers. AMPs generally interact with anionic lipid head groups, but it is less clear how the lipid tail length and saturation modulates interactions with membranes. Here we used native mass spectrometry to measure the stoichiometry of three different AMPs—LL-37, indolicidin, and magainin-2—in lipid nanodiscs. We also measured the activity of these AMPs in large unilamellar vesicle leakage assays. We found that LL-37 formed specific hexamer complexes but with different assembly pathways and affinities that depended on the bilayer thickness. LL-37 was also most active in lipid bilayers containing longer, unsaturated lipids. In contrast, indolicidin incorporated to a higher degree into more fluid lipid bilayers but was more active with thinner, less fluid bilayers. Finally, magainin-2 incorporated to a higher degree into longer, unsaturated bilayers and showed more activity in these same conditions. Together, these data show that higher amounts of peptide incorporation generally led to higher activity and that AMPs tend to incorporate more into longer unsaturated lipid bilayers. However, the activity of AMPs was not always directly related to amount of peptide incorporated.

**Graphical Abstract:** 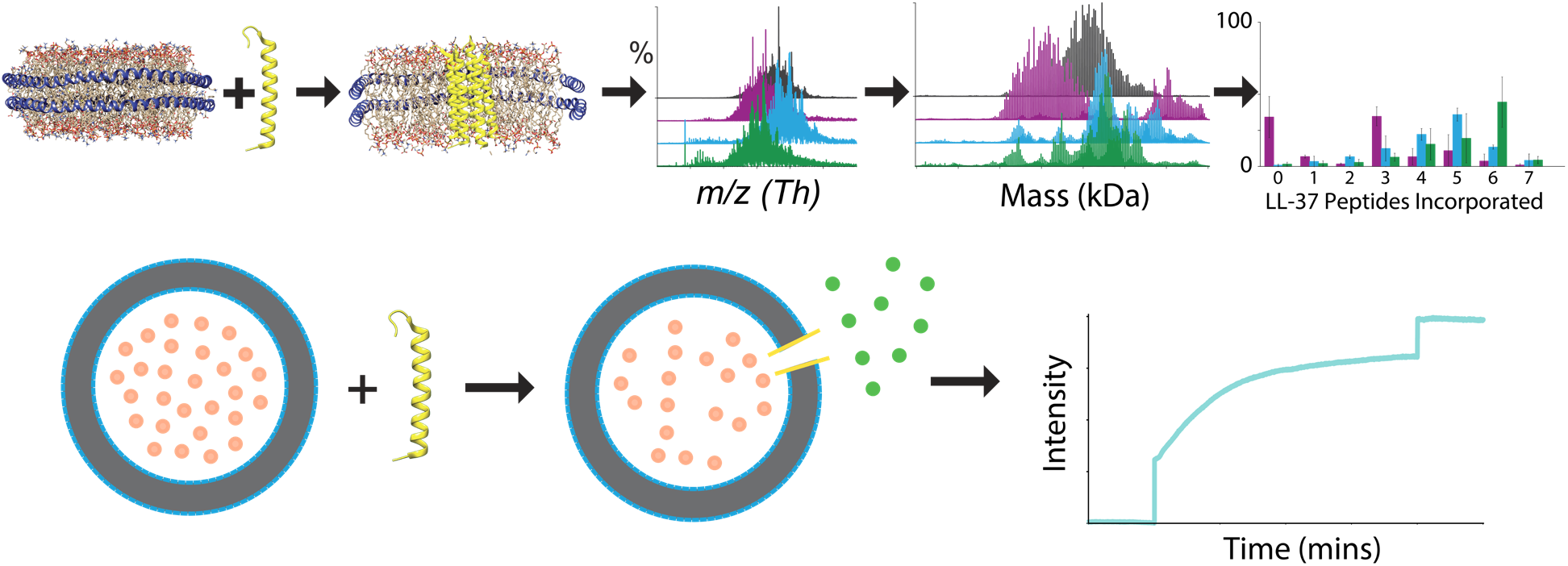

## 1. Introduction

Antimicrobial peptides (AMPs) are generally cationic, amphipathic peptides that show antimicrobial, antifungal, and antiviral activity by interacting with the lipid bilayer. AMPs are produced by a variety of organisms and act as an innate defense against infection.[1–3] However, the modes of action of AMPs are often not well understood. The human cathelicidin AMP, LL-37, has been shown to act by several different mechanisms. It may form a toroidal pore that permeabilizes the membrane,[4] cause massive disruption by globally destabilizing the cell membrane,[5] or act differently in different bilayers by forming pores in unsaturated lipid bilayers but peptide-lipid fibrils in saturated lipid bilayers.[6] Because the lipid bilayer plays a key role in each of these processes, a clearer understanding of AMP mechanisms will require a better understanding of their interactions with lipids.

Due to their cationic charge, AMPs generally have higher affinity for anionic lipid head groups present in bacterial membranes,[7–11] but it is less clear how lipid tails affect their membrane interactions. Bacteria can gain resistance to AMPs by altering both parts of lipid structure,[12] including creating less anionic head groups[13,14] and changing the membrane fluidity by modulating the fatty acids, which demonstrates that lipid tails can play an important role in AMP activity and resistance.[15,16] *In vitro* studies have shown that LL-37 insertion depth into the bilayer is dependent on lipid packing.[4,17] In phosphatidylcholine (PC) bilayers, LL-37 activity is modulated by the lipid tail length. With shorter PC lipid tails, LL-37 can form micelles, but LL-37 penetrates and interdigitates into longer tail PC bilayers.[18] Using lipid mixtures containing the same hydrophobic tail, Sevcsik et al. demonstrated that LL-37-lipid interactions are a complex interplay between head group charge and tail packing.[19] Thus, more detailed studies are needed to understand how tails of different lengths and saturations influence the incorporation of AMPs into membranes and how this incorporation correlates with AMP activity.

In previous studies, we developed a novel approach to measure the stoichiometry and specificity of AMP complexes in controlled lipid bilayer nanodiscs using native mass spectrometry (MS).[7,8] We discovered that AMPs generally incorporated more into nanodiscs containing dimyristoyl-phosphatidylglycerol (DMPG) lipids over dimyristoyl-phosphatidylcholine (DMPC), consistent with the anionic lipid head group attracting the cationic peptides.[7,8] Unlike other AMPs that were usually nonspecific, LL-37 formed specific oligomeric complexes in DMPG nanodiscs that seemed to indicate preferences for dimers, tetramers, and hexamers. Here, we explored the effect of the tail length and tail saturation on AMP incorporation into phosphatidylglycerol (PG) lipid bilayers, focusing on the specificity of LL-37 complex formation and comparing these results with indolicidin and magainin-2. We also used liposome leakage experiments to determine if the change in incorporation would cause more disruption of the lipid bilayer.

## 2. Materials and Methods

### 2.1. Materials

1,2-dilauroyl-*sn*-glycero-3-phospho-(1′-rac-glycerol) (DLPG), 1,2-dimyristoyl-*sn*-glycero-3-phospho-(1′-rac-glycerol) (DMPG), 1,2-dipalmitoyl-*sn*-glycero-3-phospho-(1′-rac-glycerol) (DPPG), 1-palmitoyl-2-oleoyl-*sn*-glycero-3-phospho-(1′-rac-glycerol) (POPG), and 1-stearoyl-2-oleoyl-*sn*-glycero-3-phospho-(1′-rac-glycerol) (SOPG) lipids were purchased from Avanti Polar Lipids (Alabaster, AL). Indolicidin, LL-37, and magainin-2 were purchased from Anaspec (Fremont, CA). All peptides were 95% or greater purity. Ammonium acetate, Amberlite XAD-2, sodium cholate, Triton X-100, and 5(6)-carboxyfluorescein were purchased from Sigma-Aldrich. Methanol (LC/MS grade) was purchased from Fischer Scientific.

### 2.2. Nanodisc Assembly

Nanodiscs were assembled as previously described.[20,21] Briefly, lipids dissolved in chloroform were dried under nitrogen and overnight under vacuum. Dried lipids were dissolved in 0.1 M sodium cholate. The lipid-cholate mixture was mixed with membrane scaffold protein, MSP1D1(–), which was purified from *E. coli* as previously described.[21,22] The lipid and MSP were mixed at a molar ratio of 100:1 for DLPG, DMPG, and DPPG lipids, and 80:1 for POPG and SOPG lipids, with a final cholate concentration of 20–25 mM. After incubating at the lipid phase transition temperature, Amberlite XAD-2 hydrophobic beads were added to the mixture to remove the cholate detergent. The nanodiscs were removed from the beads, filtered, and loaded onto a Superose 6 Increase 10/300 column to purify the nanodiscs. The running buffer was 0.2 M ammonium acetate, pH 6.8.

### 2.3. Mass Spectrometry Sample Preparation

Samples were prepared for native MS as previously described.[8] Briefly, nanodiscs were purified and diluted to 2.5 μM before being mixed with peptides at specific molar ratios. Nanodisc samples were prepared in 0.2 M ammonium acetate and 19 μL of 2.5 μM nanodiscs was mixed with 3 μL of peptide (dissolved in methanol) and 1.5 μL of 0.4 M imidazole. Peptides were mixed with nanodiscs at 3:1, 9:1, and 18:1 molar ratios of peptide:nanodisc and allowed to equilibrate for 5 minutes before measuring.

### 2.4. Native Mass Spectrometry

Native MS was performed on a Q-Exactive HF quadrupole-Orbitrap mass spectrometer with the Ultra-High Mass Range (UHMR) research modifications (Thermo Fisher Scientific, Bremen, Germany).[7] Samples were introduced into the mass spectrometer using nano-electrospray ionization with a capillary voltage of 1.1 kV and a capillary temperature of 200 °C. Samples were analyzed from 2,000–25,000 *m/z* at a resolution setting of 15,000 and a trapping gas setting of 7. Source fragmentation was set to 50 V to help aid in desolvation, and in source trapping voltage was set to 0 V.

### 2.5. Mass Spectrometry Data Analysis

Data analysis was performed as previously described.[7,8] Briefly, native mass spectra were deconvolved using UniDec and MetaUniDec.[23,24] Deconvolution settings were as follows: mass range of 20–200 kDa, charge range of 5–25, and peak full width half-maximum of 10. The lipid mass was used as the mass difference. Following deconvolution, macromolecular mass defect analysis was used to determine the number of peptides incorporated into the nanodiscs. Unfortunately, overlap between the mass of six POPG molecules and LL-37 prevented interpretation of native MS data with POPG nanodiscs (Table S1), so no results are reported for this lipid combination.

### 2.6. LUV Assembly

Large unilamellar vesicles (LUVs) were made as previously described.[25] Briefly, ~10 mg of lipid in chloroform was dried under nitrogen followed by overnight in vacuum. 500 μL of 20 mM carboxyfluorescein and 200 mM ammonium acetate, pH 6.8, were added to the dried lipids. The solution was sonicated for 30 seconds, followed by 15 seconds of vortexing. Sonicating and vortexing were repeated three times. The sample was extruded (Avanti Polar Lipids, Alabaster, AL) using a 100 nm polycarbonate Track-Etched filter (GE Healthcare). The extruded LUVs were loaded onto a HiTrap desalting column (GE Healthcare) to remove any carboxyfluorescein that was not encapsulated. The lipid content of the liposomes was quantified by measuring the total phosphorous concentration.[26,27]

### 2.7. Carboxyfluorescein Leakage Assay

Leakage assays were performed on a Horiba PTI Quantamaster 400 fluorometer using an excitation wavelength of 492 nm, emission wavelength of 515 nm, and slit width of 5 nm. To maintain the stability of DMPG liposomes, all experiments were performed at 10 °C. LUVs were first diluted to a total lipid concentration of 2 μM. 3 mL of this LUV sample was added to a cuvette and the fluorescence emission was measured for 5 mins. At 5 mins, peptide was added at a specific peptide:lipid ratio equal to the peptide:lipid ratio used for mass spectrometry measurements with nanodiscs. For example, a DMPG nanodisc contains an average of 160 lipids. To achieve the same 3:1 molar ratio as with nanodiscs we needed to add 1 peptide for every ~53 lipids in our solution. Using a lipid concentration of 2 μM, this means adding peptide to a final concentration of 38 nM to match our 3:1 peptide:nanodisc molar ratio. The number of lipids in a nanodisc varied for each lipid used in these studies and therefore, the amount of peptide added to the LUVs varied for each lipid used. After peptides were added to the LUVs, the measurement continued for 20 minutes. Finally, at 25 minutes, Triton X-100 was added to a final concentration of 0.1% to give the maximum fluorescence value. Controls with the addition of methanol alone (without dissolved AMPs) had a minimal effect on the release of carboxyfluorescein (~1-2%).

Each LUV experiment was repeated three times with a different LUV preparation. The results of each individual experiment were normalized to the maximum and minimum fluorescence values for that sample, and the mean of the normalized fluorescence was plotted with the standard error of the mean.

## 3. Results

### 3.1. Effects of lipid tails on LL-37 incorporation and activity

Our goal was to determine how bilayer thickness and lipid saturation affected LL-37 incorporation and complex formation in PG bilayers. Thus, we prepared nanodiscs containing a single PG lipid with tails of different length and degrees of saturation. LL-37 was titrated into each nanodisc, allowed to incubate, and the complex was then characterized by native MS (Figure S1A-B). We analyzed and quantified the small mass shifts upon AMP addition using mass defect analysis (Figure S1C) to determine the relative abundance of each peptide stoichiometry in nanodiscs (Figure S1D).

When LL-37 was added to nanodiscs containing the short tail lipid, DLPG, LL-37 incorporated at similar levels to DMPG nanodiscs but lost most of its specificity in complex formation (Figure 1A). LL-37 showed strong preferences for specific stoichiometries in DMPG nanodiscs but had much broader distributions of stoichiometries in DLPG nanodiscs. A truncated version of LL-37, KR-12, also lacked specificity in DMPG,[8] which indicates that the interplay between the lipid and the peptide is important for formation of specific complexes.

**Figure 1.**
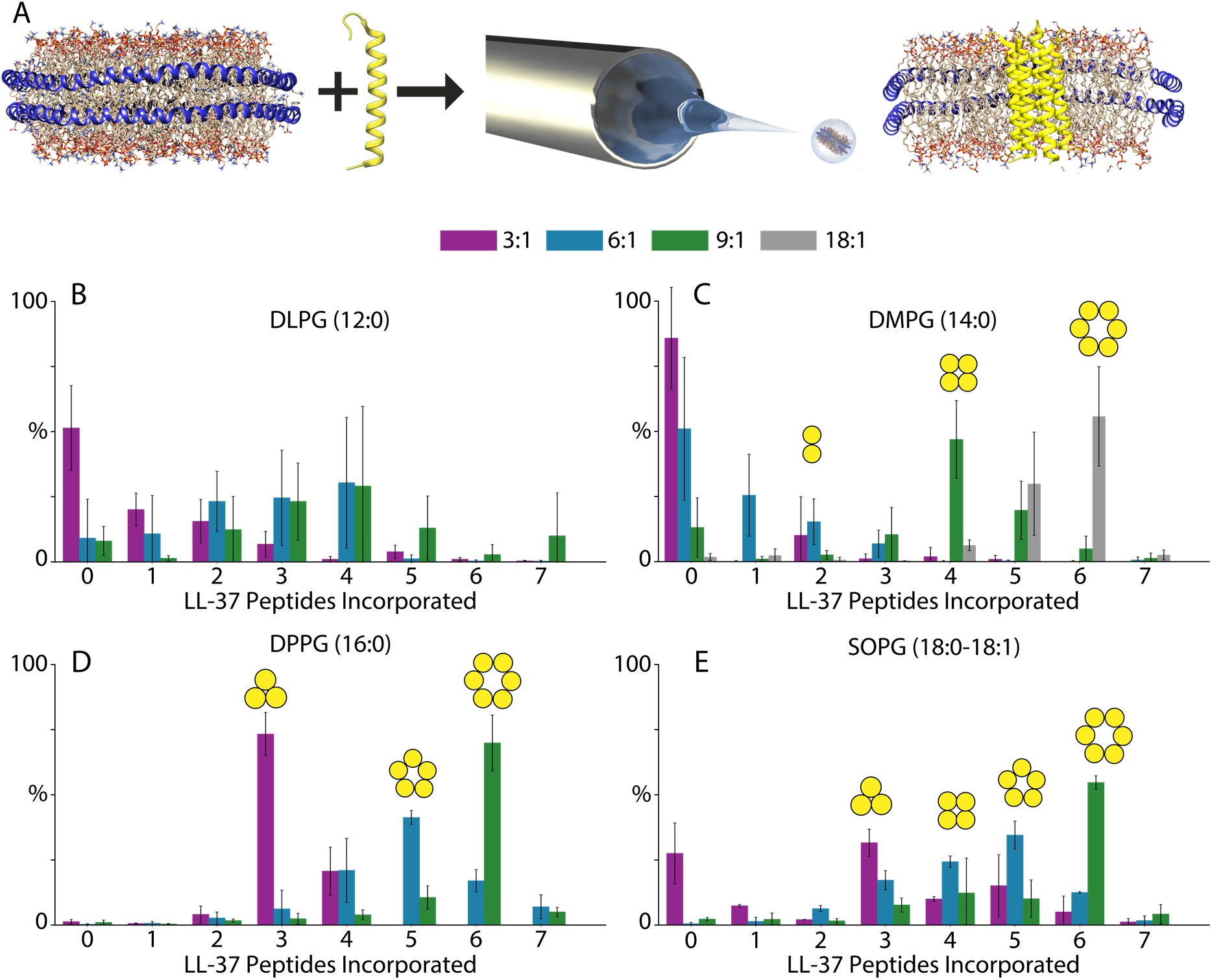
(A) Schematic of addition of LL-37 to nanodiscs. (B–E) Relative amounts of incorporation of different LL-37 stoichiometries in nanodiscs containing DLPG (A), DMPG (B), DPPG (C), or SOPG (D), at 3:1 (*purple*), 6:1 (*blue*), 9:1 (*green*), or 18:1 (*grey*) molar ratios of LL-37:nanodisc. Data for DMPG are adapted from Walker et al.[7] (C) Adapted with permission from ref. 8.

With longer lipid tails (DPPG and SOPG), the stoichiometry of LL-37 in nanodiscs was higher than observed with the shorter tail lipids at the same ratios, demonstrating an overall higher affinity for incorporation into thicker bilayers (Figure 1 and Figure 2). Higher molar ratios of LL-37 (18:1) that were stable in DMPG generally caused dissociation of the SOPG and DPPG nanodiscs. Because the highest stoichiometry observed in any lipid nanodisc was six, these data suggest that the hexamer is the highest oligomer allowed, and any further incorporation beyond this destabilizes the nanodisc, potentially through a carpet-like mechanism.

**Figure 2.**
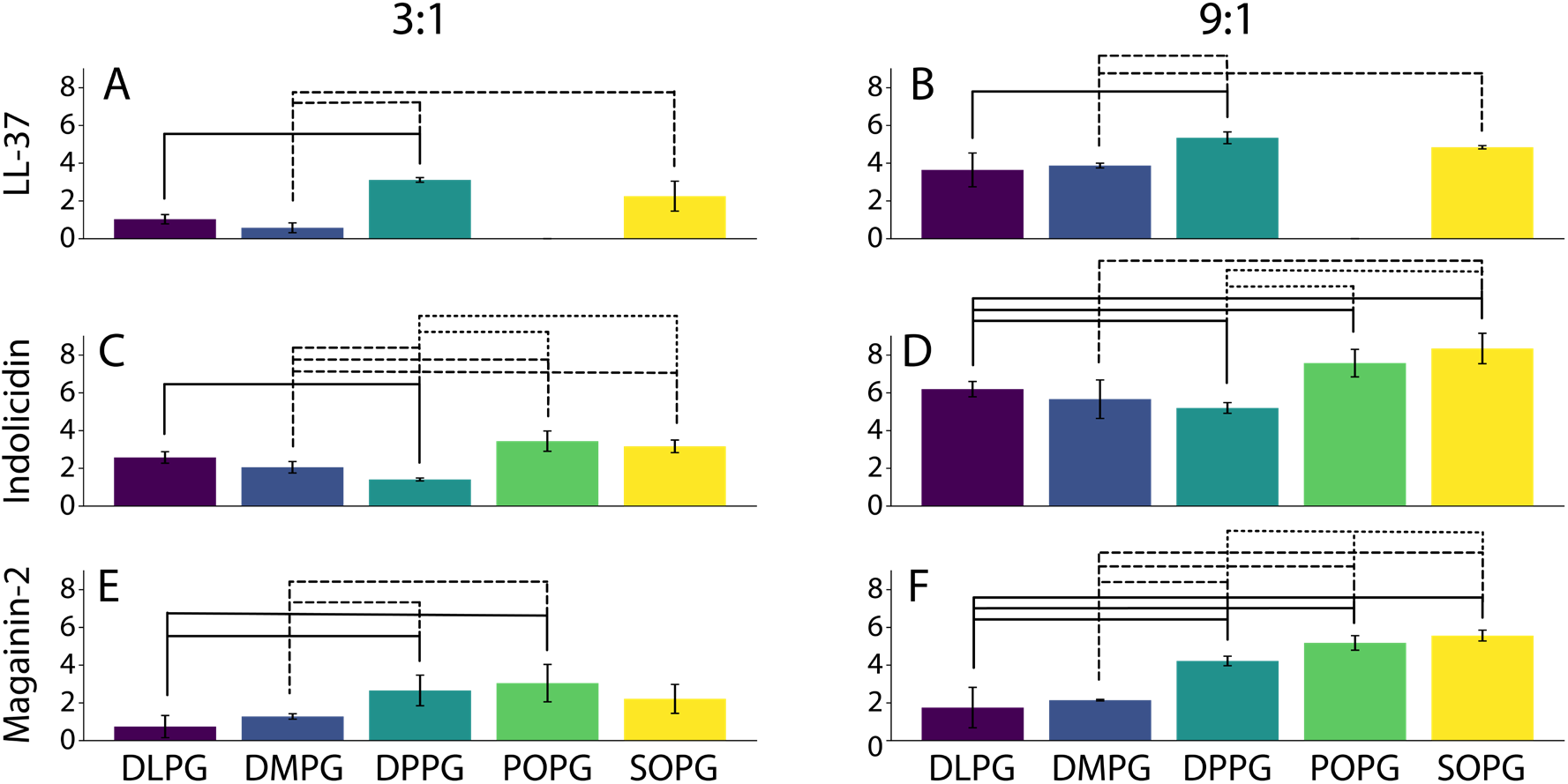
Average incorporation of peptides in nanodiscs containing different lipids at 3:1 (A, C, E) and 9:1 (B, D, F) ratios of peptide:nanodisc for LL-37 (A, B), indolicidin (C, D), and magainin-2 (E, F). Connected lines above bars indicate significance as determined by a T-test at the 95% confidence level. Data for DMPG are from Walker et al.[7]

Although native MS only reports directly on the stoichiometry of incorporation (how many peptides are associated with the nanodisc), we can infer formation of specific complexes when non-random stoichiometry distributions appear. For example, LL-37 in DMPG nanodiscs first incorporated with a stoichiometry of two, with very little nanodiscs containing one or three peptides at a 3:1 LL-37:nanodisc ratio (Figure 1C).[7] The non-random stoichiometries reveal specificity for dimer in the nanodisc. No evidence for LL-37 trimer was observed at any ratio in DMPG nanodiscs, and higher ratios show non-random distributions with a progressive increase from dimer to tetramer to hexamer, with a potential population of pentamers.[7]

In stark contrast, with DPPG and SOPG nanodiscs, trimeric LL-37 is clearly the dominant species at a 3:1 ratio. After the formation of the trimeric LL-37, additional LL-37 at higher ratios causes progressively higher stoichiometries before ultimately settling on hexamer complexes. Higher stoichiometries beyond six were not observed, demonstrating the specific formation of hexamer complexes. It is not clear whether tetramer and pentamer intermediates are specific, but there is a clear specificity for trimer and hexamer.

Together, these data reveal a fascinating picture of how lipid tails modulate formation of LL-37 complexes in PG bilayers. Overall, LL-37 has a higher affinity for thicker membranes. The similar stoichiometries between DPPG and SOPG, which have similar bilayer thickness[28] but different fluidity, suggest that membrane fluidity and unsaturation are less important than bilayer thickness. Very thin DLPG bilayers largely disrupt the formation of specific complexes. DMPG, DPPG, and SOPG all showed clear evidence for formation of specific hexamers, but the assembly pathways diverged between them. DMPG caused formation of hexamers via dimer intermediates whereas DPPG and SOPG both drove hexamer formation through trimer intermediates. Thus, lipid tails play clear roles in modulating the overall membrane affinity, the specificity of complex formation, and the intermediate building blocks used for complex assembly.

### 3.2. Effects of lipid tails on LL-37 activity

We also tested the effects of lipid tails on the activity of LL-37 using a vesicle leakage assay with carboxyfluorescein-encapsulated LUVs made with either DMPG or SOPG.[25] Although we could not match the concentrations used for native MS with nanodiscs, we titrated AMPs at similar peptide:lipid ratios with LUVs. Release of carboxyfluorescein was determined by an increase in fluorescence due to lower self-quenching upon dilution as it escaped the LUV.

When LL-37 was added to LUVs at LL-37:lipid molar ratios equivalent to 3:1, 9:1, and 18:1 LL-37:nanodisc, SOPG LUVs showed significantly more leakage of carboxyfluorescein compared to LUVs containing DMPG (Figure 3). SOPG LUVs showed partial vesicle leakage at 3:1 and nearly complete leakage (compared to addition of Triton X-100) at 9:1 and 18:1 ratios. These results agree with native MS data that showed formation of the hexamer complex at a 9:1 ratio and dissociation of nanodiscs beyond this point.

**Figure 3.**
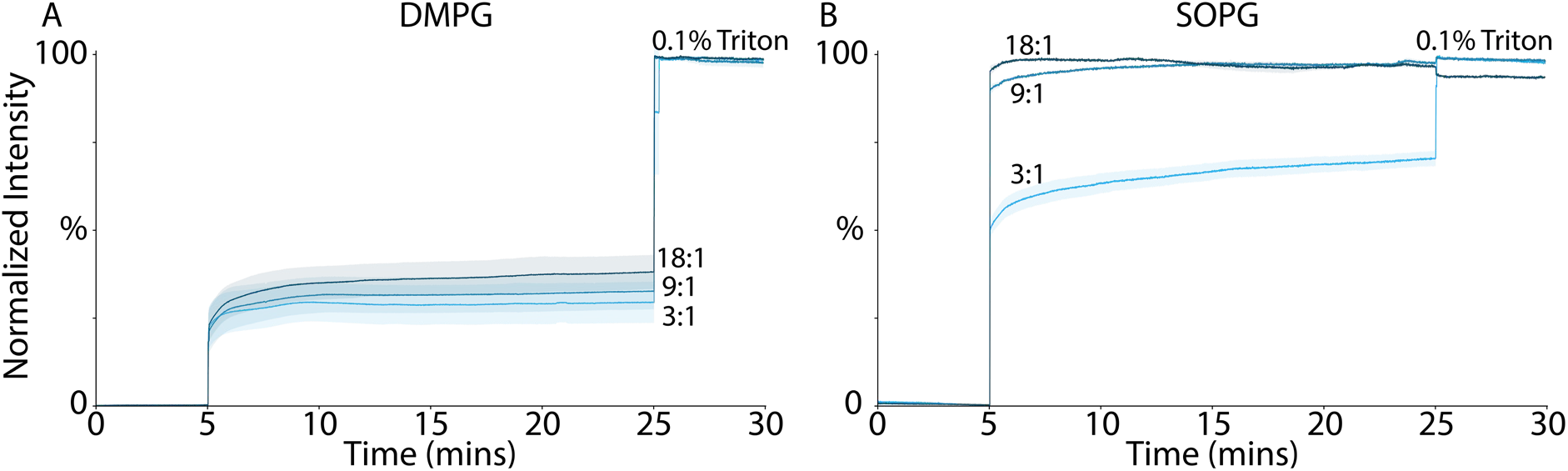
LUV Leakage Assay with DMPG (A) or SOPG (B) LUVs with addition of 3:1, 9:1, or 18:1 ratios of LL-37:nanodisc. Fluorescence of carboxyfluorescein encapsulated LUVs was measured for 5 minutes. At 5 minutes, LL-37 was added at the molar ratios indicated. At 25 minutes, 0.1% Triton was added to determine maximum fluorescence. Results were normalized to highest and lowest fluorescence values, and the mean of the normalized fluorescence from 3 different LUV preparations was calculated. The shaded areas indicate the standard error of the mean for each stoichiometry.

DMPG LUVs also showed partial vesicle leakage but less than SOPG LUVs, confirming the higher activities of LL-37 in SOPG bilayers. However, despite native MS evidence for a hexamer complex in both conditions, less leakage was observed in 18:1 DMPG compared to 9:1 SOPG. This lower activity could be due to differences in absolute concentration or temperature between the nanodisc and LUV experiments or because hexamer complexes in DMPG are not structurally identical to hexamers in SOPG, potentially forming smaller pores incapable of leaking carboxyfluorescein. In any case, the higher activity of LL-37 in SOPG over DMPG indicates that lipid tails can modulate AMP activity, with higher activity shown for longer tail lipids.

### 3.3. Effects of lipid tails on indolicidin and magainin-2 incorporation and activity

To place our results with LL-37 in context, we compared them with similar data from two different peptides, indolicidin and magainin-2. When indolicidin was titrated into PG nanodiscs containing different lipid tails, more indolicidin was incorporated into nanodiscs containing an unsaturated lipid (POPG and SOPG), with saturated lipids all showing similar amounts of incorporation (Figure 2C-D, S2, and S3). Interestingly, indolicidin showed the lowest level of incorporation in nanodiscs containing DPPG. This indicates that indolicidin incorporation is not controlled by bilayer thickness but instead by bilayer saturation/fluidity.

We then compared the native MS nanodisc data to the LUV leakage assay. At low molar ratios (3:1), indolicidin caused significantly more leakage in DMPG LUVs than SOPG LUVs (Figure S6). This contrasted the higher stoichiometries observed in SOPG nanodiscs over DMPG nanodiscs (Figure 2C). However, indolicidin formed tighter distributions in DMPG nanodiscs that may indicate more specific oligomeric complexes rather than nonspecific membrane association (Figure S2). As described above, it may be that differences in temperature, concentration, or other experimental parameters cause this apparent discrepancy between incorporation stoichiometries measured by native MS and activity measured by LUV leakage. It could also be that the more specific indolicidin complexes in DMPG are more active than the less specific complexes in SOPG. At an intermediate stoichiometry (9:1), indolicidin showed similar levels of carboxyfluorescein leakage after 20 minutes. However, SOPG LUVs took longer to reach the same level of leakage while the leakage from DMPG LUVs was almost instantaneous (Figure S6). At the highest stoichiometry tested (18:1), there was fast and complete leakage in both DMPG and SOPG LUVs (Figure S6). Overall, indolicidin incorporated more into more fluid membranes but was more active in less fluid membranes, potentially by forming more specific complexes.

Like indolicidin, we previously found that the AMP magainin-2 showed minimal specificity in DMPG nanodiscs.[8] When different lipid tails were used, magainin-2 incorporated more into longer and more unsaturated tails (Figure 2E-F, Figure S4 & S5). At a 9:1 ratio, the highest incorporation was into nanodiscs containing SOPG, followed by POPG, and then DPPG. This data indicates that magainin-2 prefers both longer and more unsaturated lipid tails. Like indolicidin, magainin-2 generally showed little preference for specific complex formation (Figures S4 & S5).

The SOPG LUVs showed ~100% carboxyfluorescein leakage after 20 minutes at all molar ratios of magainin-2 tested but steadily increasing rates at higher concentrations (Figure S7B). With DMPG LUVs, the amount of carboxyfluorescein leakage also increased with concentration but never went higher than ~50% leakage after 20 minutes (Figure S7A). For this peptide, the LUV leakage data shows a strong correlation with the incorporation data, indicating that magainin-2 has higher activity and higher incorporation in longer and more unsaturated lipid bilayers.

## 4. Conclusions

Here, we investigated how PG lipid tails affect the incorporation of the AMP LL-37 into lipid nanodiscs and how this incorporation affects the leakage of carboxyfluorescein from LUVs. Overall, LL-37 showed higher levels of incorporation and activity in lipid bilayers containing longer lipids but was relatively unaffected by unsaturation. Interestingly, LL-37 formed specific hexamer complexes in all but DLPG but showed a dimer-mediated assembly pathway in DMPG and a trimer-mediated assembly pathway in DPPG and SOPG, showing that the lipid tails can also influence oligomeric assembly pathways.

We also investigated two other AMPS, indolicidin and magainin-2. Indolicidin incorporated more in more fluid membranes but had higher leakage activity in less fluid membranes. In contrast, magainin-2 showed both higher incorporation and a higher level of vesicle leakage with longer, unsaturated lipid tails. Together, these results reveal the intricate roles of lipid tails in AMP interactions with membranes, affecting the overall membrane affinity, the assembly of specific complexes, and the ability to disrupt lipid bilayers.

## Acknowledgments

This work was funded by the National Institute of General Medical Sciences and the National Institutes of Health (R35 GM128624). The authors thank Maria Reinhardt-Szyba, Kyle Fort, and Alexander Makarov at Thermo Fisher Scientific for support with the Q-Exactive HF instrument. The pMSP1D1 plasmid was a gift from Stephen Sligar (Addgene plasmid number 20061). The content is solely the responsibility of the authors and does not necessarily represent the official views of the NIH.

