## Supporting Information for "Lipid Tails Modulate Antimicrobial Peptide Membrane Incorporation and Activity"

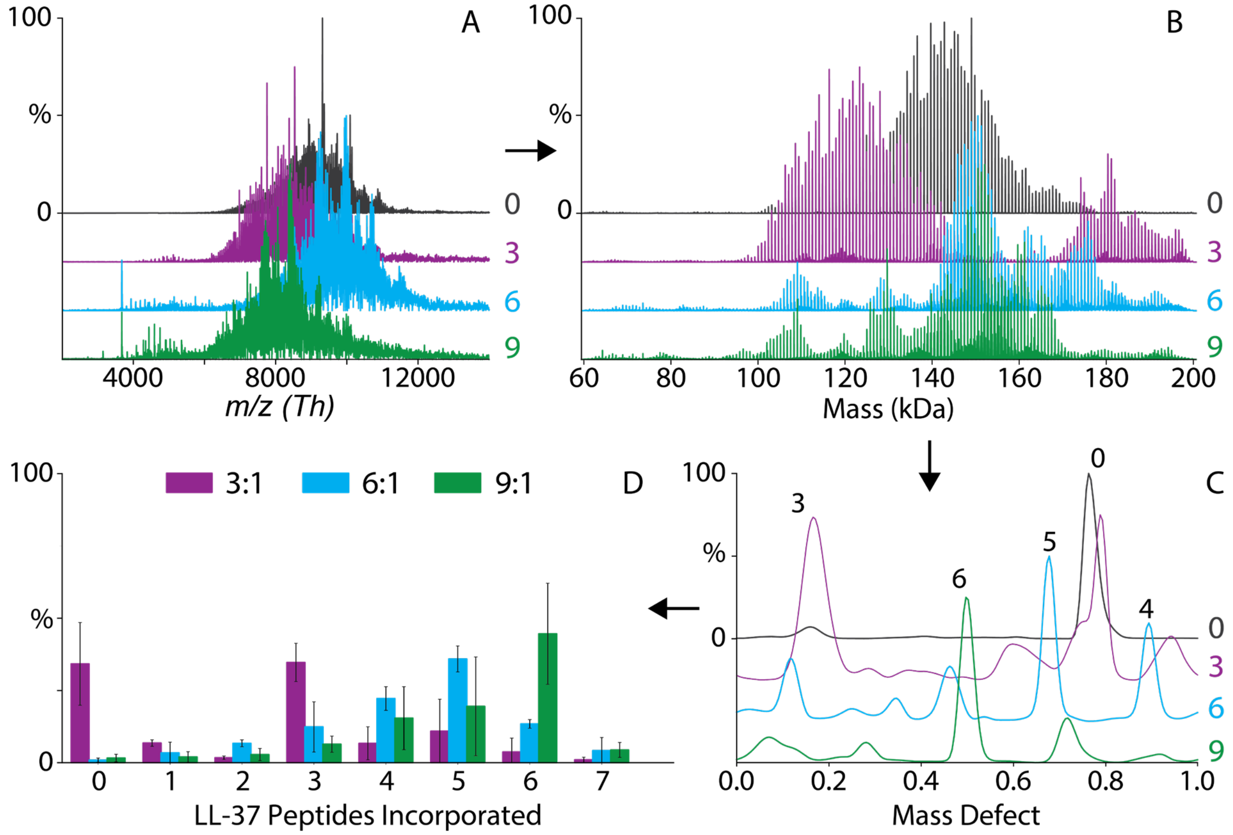


**Figure S1.** LL-37 incorporation into SOPG nanodiscs. A) Raw mass spectra of of nanodiscs with LL-37 from 0 to 9:1 molar ratio of LL-37 to nanodiscs. B) Deconvolved mass spectra. C) Mass defect analysis showing different stoichiometries of LL-37 incorporated. D) Relative amounts of LL-37 incorporation into SOPG nanodiscs.

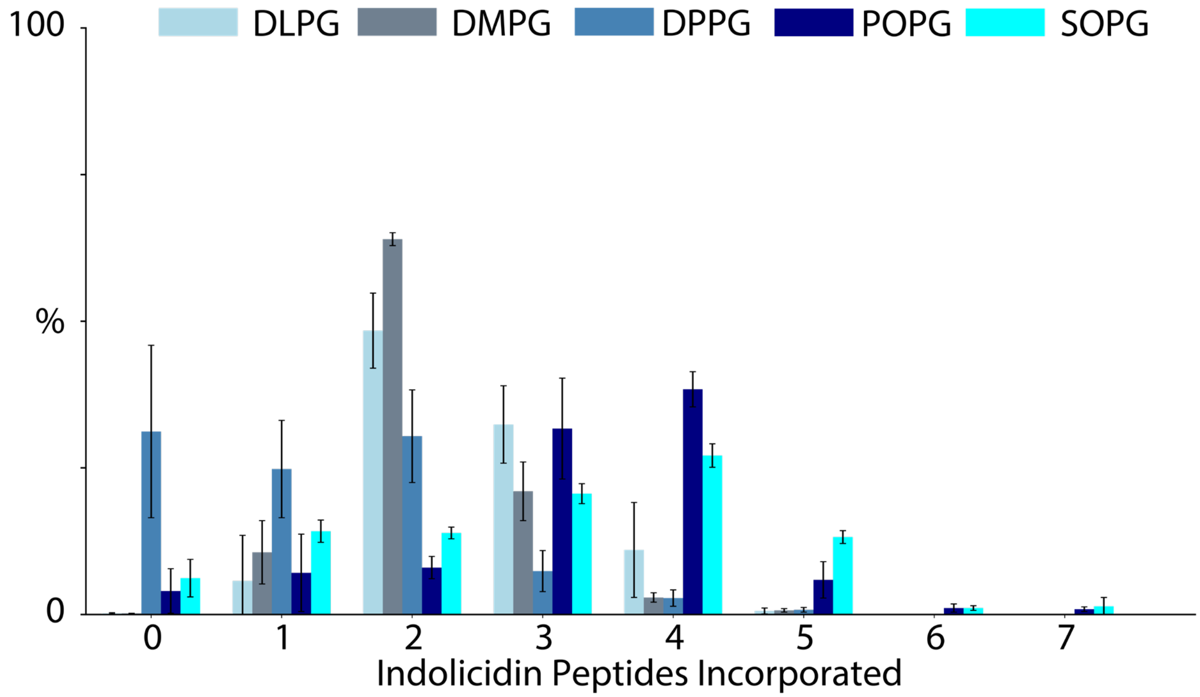


**Figure S2.** Indolicidin incorporation in nanodiscs at a 3:1 molar ratio of indolicidin to nanodiscs with different PG lipids (shown in different colors).

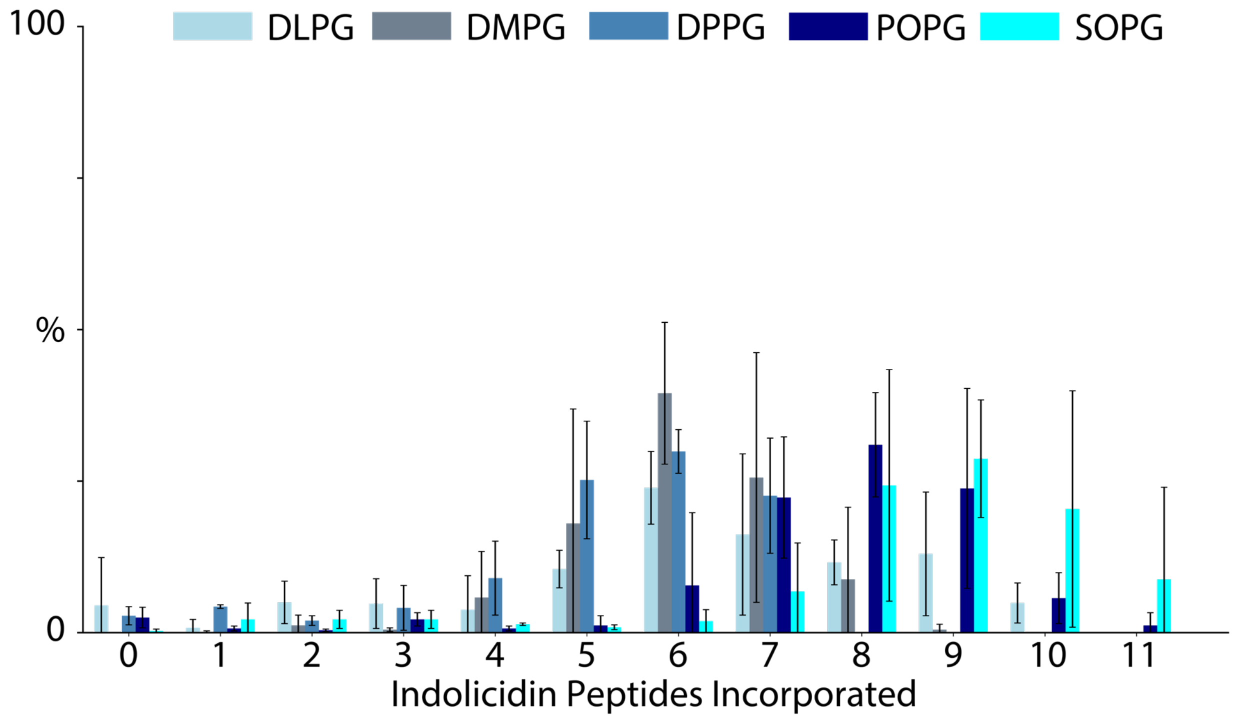


**Figure S3.** Indolicidin incorporation in nanodiscs at a 9:1 molar ratio of Indolicidin to nanodiscs with different PG lipids (shown in different colors).

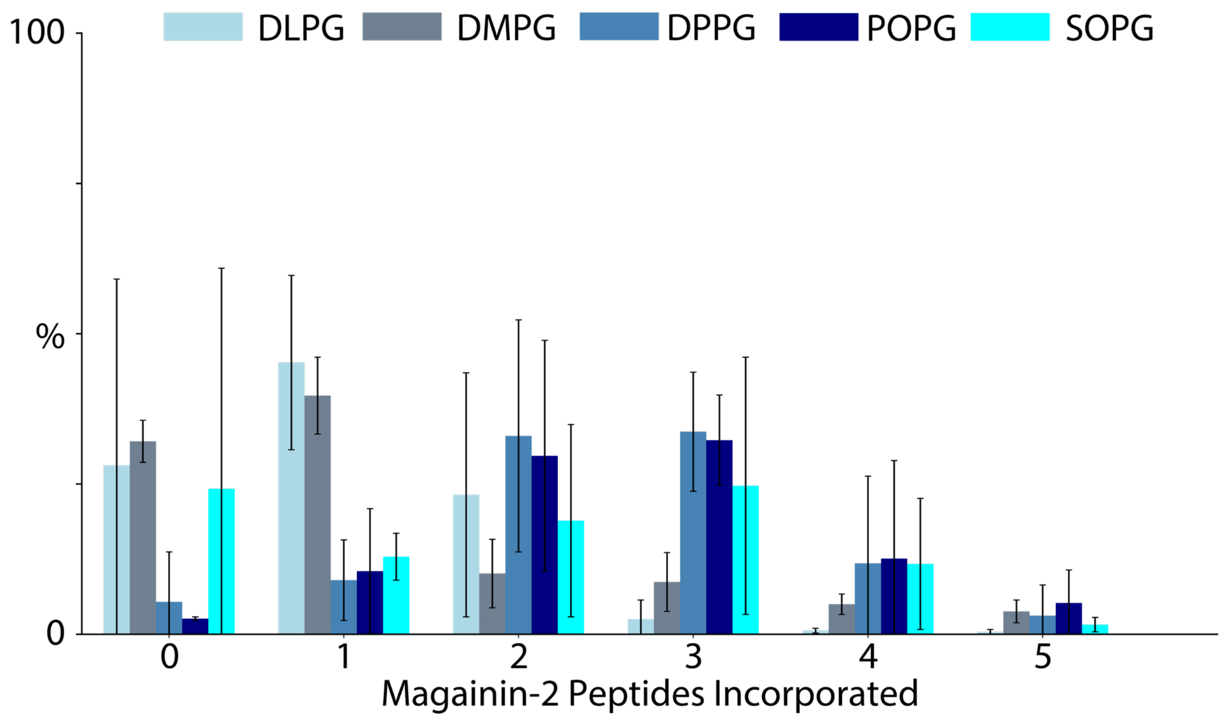


**Figure S4.** Magainin-2 incorporation in nanodiscs at a 3:1 molar ratio of magainin-2 to nanodiscs with different PG lipids (shown in different colors).

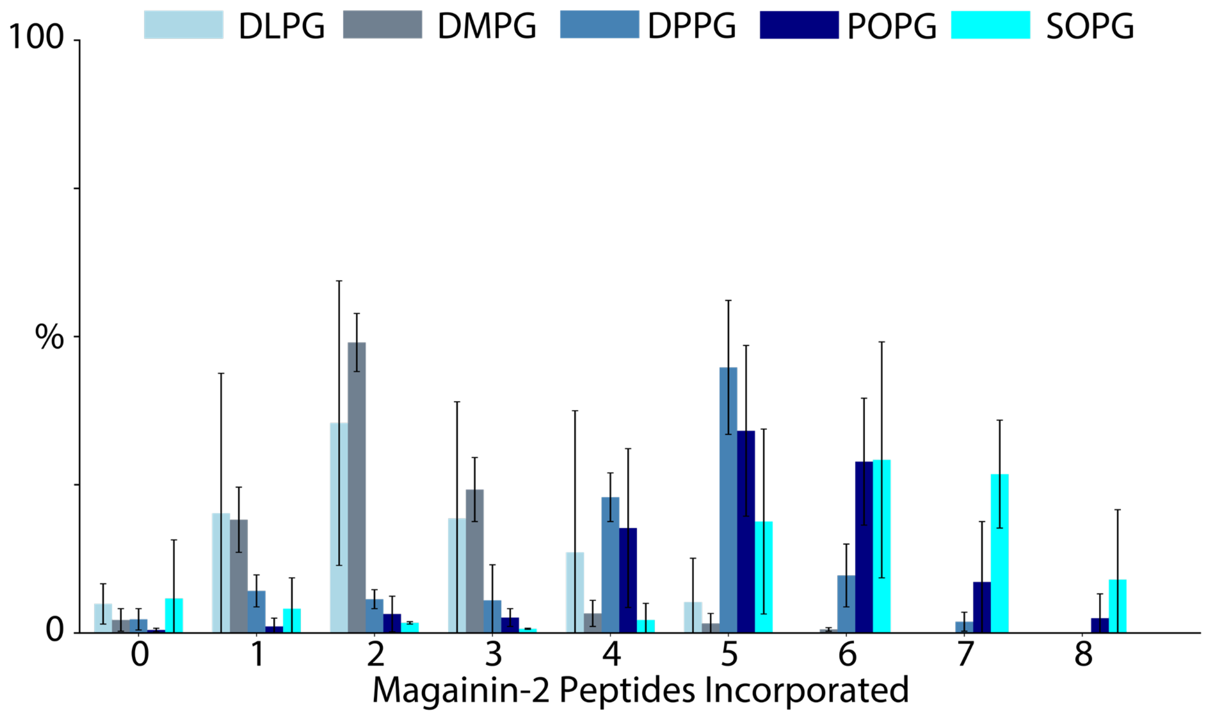


**Figure S5.** Magainin-2 incorporation in nanodiscs at a 9:1 molar ratio of magainin-2 to nanodiscs with different PG lipids (shown in different colors).

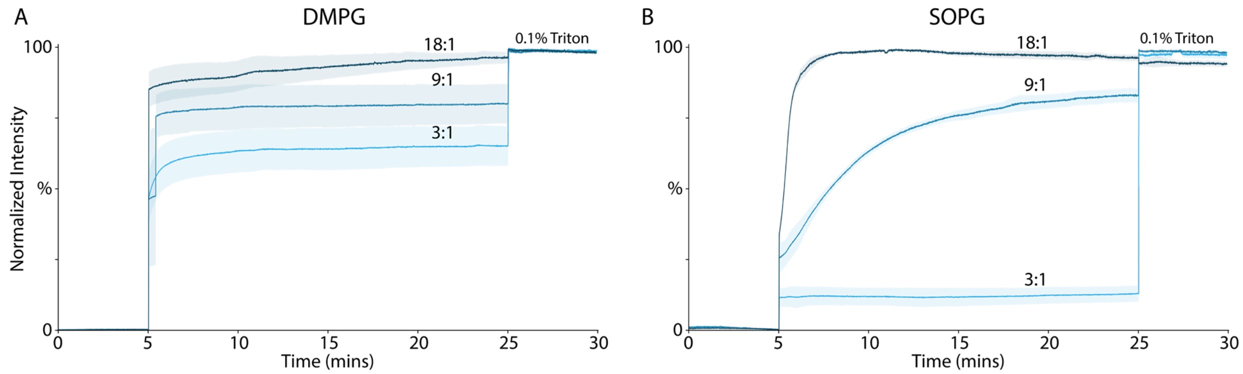


**Figure S6.** LUV fluorescence of DMPG (A) or SOPG (B) LUVs with 3:1, 9:1, or 18:1 indolicidin. Fluorescence of carboxyfluorescein encapsulated LUVs was measured for 5 minutes. At 5 minutes, peptide was added at specific molar ratios indicated. At 25 minutes, 0.1% Triton was added to determine maximum fluorescence. Results were normalized to highest and lowest fluorescence values, and the mean of the normalized fluorescence from 3 different LUV preparations was calculated. The shaded areas indicate the standard error of the mean for each stoichiometry.

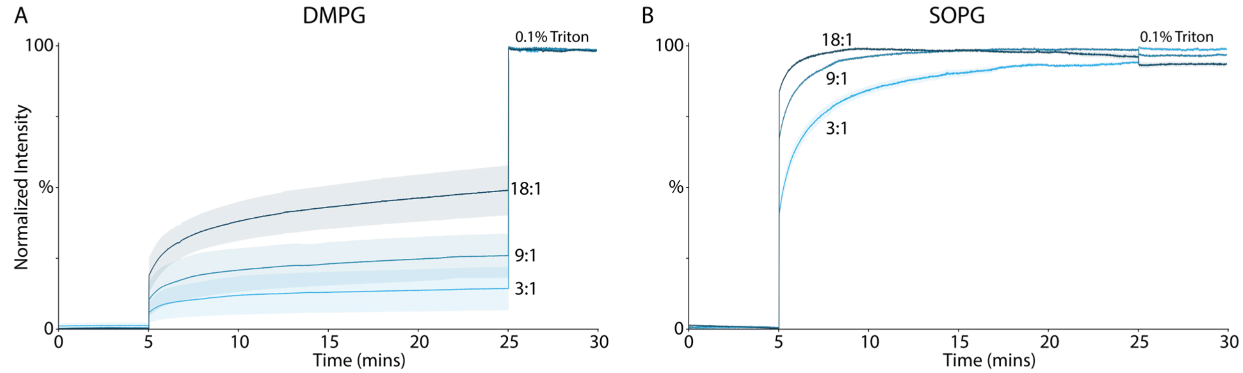


**Figure S7.** LUV fluorescence of DMPG (A) or SOPG (B) LUVs with 3:1, 9:1, or 18:1 magainin-2. Fluorescence of carboxyfluorescein encapsulated LUVs was measured for 5 minutes. At 5 minutes, magainin-2 was added at specific molar ratios indicated. At 25 minutes, 0.1% Triton was added to determine maximum fluorescence. Results were normalized to highest and lowest fluorescence values, and the mean of the normalized fluorescence from 3 different LUV preparations was calculated. The shaded areas indicate the standard error of the mean for each stoichiometry.

Table S1: Antimicrobial peptides used in this study.

| Peptide | Sequence | Charge | Mass (Da) | Mass Defect Values | | | | |
|---|---|---|---|---|---|---|---|---|
| | | | | DLPG | DMPG | DPPG | POPG | SOPG |
| Indolicidin | ILPWKWPWWPWRR | +3 | 1906.3 | 0 - 0.19 | 0.10 | 0.98 | 0.86 | 0.74 |
| | | | | 1 - 0.31 | 0.96 | 0.62 | 0.41 | 0.19 |
| | | | | 2 - 0.44 | 0.81 | 0.25 | 0.95 | 0.65 |
| | | | | 3 - 0.56 | 0.67 | 0.89 | 0.50 | 0.10 |
| | | | | 4 - 0.68 | 0.53 | 0.53 | 0.04 | 0.55 |
| | | | | 5 - 0.80 | 0.39 | 0.16 | 0.59 | 0.01 |
| | | | | 6 - 0.92 | 0.25 | 0.80 | 0.13 | 0.46 |
| | | | | 7 - 0.04 | 0.11 | 0.44 | 0.68 | 0.92 |
| | | | | 8 - 0.16 | 0.96 | 0.07 | 0.22 | 0.37 |
| | | | | 9 - 0.29 | 0.82 | 0.71 | 0.77 | 0.82 |
| | | | | 10 - 0.41 | 0.68 | 0.35 | 0.31 | 0.28 |
| | | | | 11 - 0.53 | 0.54 | 0.98 | 0.86 | 0.73 |
| LL-37 | LLGDFFRKSKEKIGKEFKRIVQRIKDFLRNLVPRTES | +6 | 4493.3 | 0 - 0.19 | 0.10 | 0.98 | 0.86 | 0.74 |
| | | | | 1 - 0.55 | 0.84 | 0.19 | 0.86 | 0.52 |
| | | | | 2 - 0.91 | 0.57 | 0.41 | 0.86 | 0.31 |
| | | | | 3 - 0.27 | 0.31 | 0.62 | 0.86 | 0.09 |
| | | | | 4 - 0.62 | 0.05 | 0.84 | 0.86 | 0.87 |
| | | | | 5 - 0.98 | 0.78 | 0.05 | 0.86 | 0.66 |
| | | | | 6 - 0.34 | 0.52 | 0.27 | 0.86 | 0.44 |
| | | | | 7 - 0.70 | 0.26 | 0.48 | 0.86 | 0.22 |
| Magainin-2 | GIGKFLHSAKKFGKAFVGEIMNS | +4 | 2467.1 | 0 - 0.19 | 0.10 | 0.98 | 0.86 | 0.74 |
| | | | | 1 - 0.23 | 0.80 | 0.39 | 0.16 | 0.92 |
| | | | | 2 - 0.27 | 0.50 | 0.80 | 0.45 | 0.09 |
| | | | | 3 - 0.31 | 0.19 | 0.22 | 0.74 | 0.27 |
| | | | | 4 - 0.35 | 0.89 | 0.63 | 0.04 | 0.44 |
| | | | | 5 - 0.39 | 0.59 | 0.04 | 0.33 | 0.62 |
| | | | | 6 - 0.43 | 0.29 | 0.45 | 0.62 | 0.79 |
| | | | | 7 - 0.47 | 0.99 | 0.86 | 0.92 | 0.97 |
| | | | | 8 - 0.51 | 0.69 | 0.28 | 0.21 | 0.14 |
